# In-cell structural analysis reveals a distinctive chloroplast ribosome in *Chlamydomonas reinhardtii*

**DOI:** 10.64898/2026.08.28.747797

**Authors:** Zhen Hou, Yao Shen, Zhaobo Zhang, Pengchi Lu, Aris Katzourakis, Peijun Zhang

## Abstract

Chloroplast ribosomes synthesize plastid-encoded components of photosynthetic machinery, yet their structure and organization remain poorly understood. We combined cryo-focused ion beam milling, cryo-electron tomography and subtomogram averaging to determine native chloroplast ribosomes in *Chlamydomonas reinhardtii*. The 4.4–4.9 Å structure revealed a large arch-like extension on the small subunit (SSU). Comparisons with bacterial and plant chloroplast ribosomes, supported by proteomics, AlphaFold3 predictions and a recent atomic model, indicate that the arch is formed by insertions and extensions in SSU proteins. Classification resolved active, thylakoid-associated ribosomes with density adjacent to the nascent peptide exit and an arch-moved state enriched among thylakoid-associated particles, with coordinated displacement of the arch and beak. Phylogenetic analysis revealed an evolutionary mosaic: the uS3c insertion is broadly distributed across Chlorophyceae, whereas the uS2c insertion, uS5c and PSRP7 are concentrated in Chlamydomonadales, with PSRP7 also in Sphaeropleales. Nuclear-encoded components were recruited stepwise onto a plastid-encoded scaffold, with all four under comparable purifying selection. These findings link a lineage-specific SSU extension to ribosome dynamics, thylakoid association and evolution, highlighting the value of in-cell structural analysis.

## Introduction

Chloroplasts arose from the endosymbiotic incorporation of a cyanobacterial ancestor and retain a reduced genome together with the machinery required for its expression^1–3^. Although most ancestral genes have been transferred to the nucleus, chloroplast genomes continue to encode core subunits of photosystem I, photosystem II, the cytochrome *b6f* complex, ATP synthase and ribulose-1,5-bisphosphate carboxylase (RuBisCo), as well as components of the organellar gene-expression machinery^1–4^. These proteins are synthesized by bacterial-type 70S chloroplast ribosomes and assembled with nucleus-encoded subunits imported from the cytosol. Chloroplast translation is therefore essential for the biogenesis, maintenance and repair of the photosynthetic apparatus and provides a major regulatory point through which protein production is coordinated with development, metabolism and environmental conditions^5–7^. The unicellular green alga *Chlamydomonas reinhardtii* has been particularly valuable for dissecting these processes because its chloroplast genome, genetics and photosynthetic physiology are experimentally accessible^8,9^.

Chloroplast ribosomes retain the conserved bacterial ribosomal core but have acquired lineage-specific proteins, insertions and terminal extensions during plastid evolution. Proteomic analyses of spinach chloroplast ribosomes established the composition of the 30S and 50S subunits and identified a set of plastid-specific ribosomal proteins, several of which compensate for changes in ribosomal RNA or form additional surfaces implicated in translational regulation^10,11^. Corresponding analyses in *C. reinhardtii* revealed even more extensive divergence. In particular, the small subunit (SSU) contains unusually large homologues of the bacterial proteins S2, S3 and S5, as well as a distinctive S1-domain-containing protein PSRP7 and additional ribosome-associated factors^12,13^. Broader chloroplast proteomic and ribosome-interactome studies further indicated that chloroplast ribosomes associate with numerous proteins involved in RNA metabolism, translation regulation, redox control and photosystem biogenesis^14,15^. Together with examples of protein replacement during chloroplast ribosome evolution^16^, these findings showed that the plastid translation machinery is more compositionally diverse than its conserved bacterial core might suggest.

Structural studies of purified spinach chloroplast ribosomes subsequently established the molecular basis of several land-plant-specific features^17,18^. High-resolution cryo-electron microscopy (cryo-EM) reconstructions defined modifications of the large subunit, the peptide-exit region, the mRNA entry and exit channels and the locations of plastid-specific proteins on the SSU^17–20^. Structures containing the chloroplast ribosome-recycling factor and hibernation-promoting factor further demonstrated how chloroplast-specific components regulate ribosome activity^21^. Genetic and biochemical studies have also shown that some plastid-specific proteins are integral to ribosome assembly, whereas others function as transient regulatory factors; notably, PSRP1 is a homologue of the bacterial hibernation factor pY rather than a constitutive ribosomal protein^22,23^. However, these structural studies were almost exclusively based on purified ribosomes from flowering plants. Although one early cryo-EM study tried to resolve the structure of the *C. reinhardtii* chloroplast ribosome, the resulting reconstruction did not show any prominent feature corresponding to the pronounced enlargement of the *C. reinhardtii* SSU inferred from proteomics^24^. Therefore, this puzzle remained without a clear structural explanation for more than two decades^13^.

In living cells, chloroplast translation is linked to the spatial organization of chloroplast ribosomes, the abundance of free and thylakoid-bound ribosomes varies with the cell cycle and illumination, and ribosomes and specific mRNAs concentrate at membrane-associated sites involved in photosystem biogenesis^25–31^. However, these biochemical and imaging approaches cannot resolve the conformational states or local organization of individual ribosomes. Cryo-focused ion beam (cryo-FIB) milling and cryo-electron tomography (cryo-ET) now enable macromolecular structures and functional states to be examined directly in vitrified cells^32–39^. A recent study applying these methods to *C. reinhardtii* identified a large SSU extension domain formed predominantly by extensions and insertions in conserved ribosomal proteins and showed that a subset of ribosomes is flexibly associated with thylakoid membranes^40^. Nevertheless, how movement of this lineage-specific domain relates to membrane association, whether membrane-bound and stromal ribosomes differ in their spatial organization, and how the underlying protein extensions are distributed and constrained during evolution remained unresolved.

Here, we combine cryo-FIB milling, cryo-ET and subtomogram averaging (STA) to determine the native structure and organization of *C. reinhardtii* chloroplast ribosomes in cells. Our 4.9 Å reconstruction reveals a prominent arch-like SSU extension formed mainly by insertions and extensions in conserved SSU proteins. Classification identifies a thylakoid-associated population with connecting density adjacent to the nascent peptide exit and resolves an arch-moved state coupled to displacement of the SSU beak and strongly enriched among membrane-bound particles. Finally, phylogenetic analyses show that the chloroplast ribosome arch evolved through the stepwise recruitment of younger, nuclear-encoded components onto an older plastid-encoded scaffold, with all major components subsequently maintained by purifying selection. Together, these findings connect a lineage-specific ribosome structure to conformational heterogeneity, thylakoid association and spatial organization, and illustrate how in-cell structural analysis can uncover molecular relationships that remain inaccessible to studies of purified complexes.

## Results

### In-cell structure of native chloroplast ribosomes resolved by cryo-ET

To resolve the structure and visualize the organization of native chloroplast ribosomes in their cellular context, we prepared cryo-FIB-milled lamellae from *C. reinhardtii* cells and collected cryo-ET tilt series from chloroplast-containing regions. The reconstructed tomograms resolved the chloroplast stroma, thylakoid membranes and adjacent cellular compartments (Fig. 1a, b and Supplementary Video 1). We initially identified chloroplast ribosomes by template matching using a low-pass-filtered spinach chloroplast ribosome map (EMD-3533) as the reference^17^. Chloroplast ribosomes could be distinguished from cytoplasmic ribosomes based on their location during particle picking and segmentation (Fig. 1b and Supplementary Fig. 1a). Mapping the particles back into the tomograms revealed chloroplast ribosomes distributed throughout the stroma, frequently occupying the spaces between neighbouring thylakoid membranes. Following manual inspection and removal of false positives, an initial reconstruction at approximately 16 Å resolution revealed additional density that substantially increased the overall dimensions of the particle relative to the reference structure (Fig. 1b, enlarged view, and Supplementary Fig. 1a).

**Figure 1.**
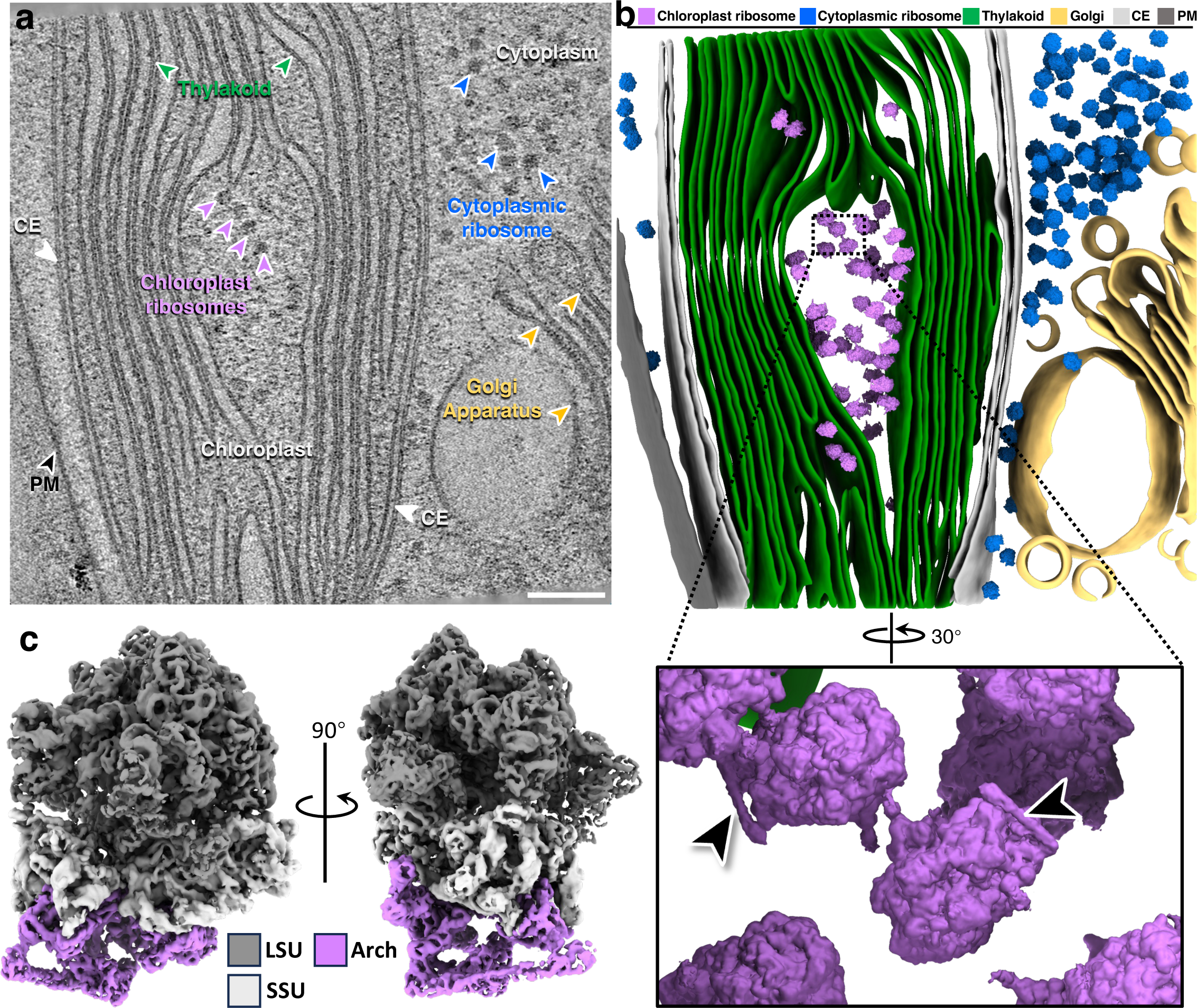
Cryo-ET of the chloroplast and in-cell structure of native chloroplast ribosomes in *C. reinhardtii*. **a,** A representative tomographic slice of the chloroplast. Chloroplast ribosomes, cytosolic ribosomes, thylakoid, Golgi apparatus (Golgi), chloroplast envelope (CE), and plasma membrane (PM) are labelled. The chloroplast and and cytoplasm are annotated accordingly. Scale bar, 100 nm. **b**, The segmented volume of (**a**), shown as an overview (top) and zoomed-in views of chloroplast ribosomes (bottom). Chloroplast ribosomes, cytoplasmic ribosomes, thylakoid, Golgi, CE (chloroplast envelope), and PM (plasma membrane) are mapped back and segmented with indicated colours. The extra density on small subunits are indicated by arrowheads, the model is the in-cell structure resolved at 16 Å (binning of 4×) in this study. **c**, Subtomogram average (STA) consensus EM density map of the chloroplast ribosome at 4.9 Å resolution (binning of 1×). The large subunit (LSU), small subunit (SSU), and extra density (the “Arch”) are coloured and annotated accordingly. Two orthogonal views are displayed.

We therefore performed iterative 3D classification and refinement to define this distinguishing structural feature. Refinement of 34,934 particles yielded a consensus map at 4.9 Å resolution, with local resolution reaching 4.4 Å and secondary-structure elements resolved in the best-defined regions (Fig. 1c, Supplementary Fig. 1a, b, d, e and Supplementary Table 3). The resulting in-cell structure comprises the conserved large subunit (LSU) and small subunit (SSU), together with a prominent arch-like density extending from the SSU, which we hereafter refer to as the arch (Fig. 1c).

### SSU-protein insertions and extensions form the arch

To define the composition and position of the arch, we compared our in-cell STA map with the *E. coli* ribosome and the *S. oleracea* chloroplast ribosome (Fig. 2a). The arch extends the long-axis dimension of the particle from approximately 220 Å to 295 Å, spanning the platform, body, and beak of the SSU near the mRNA entry and exit channels (Supplementary Fig. 2a). Apart from this region, the SSU and LSU closely resemble the conserved bacterial and plant chloroplast ribosome cores. Accordingly, rigid-body docking of the orthologous structures accounted for most ribosomal proteins and rRNA but left the arch-shaped density unexplained (Fig. 2b).

**Figure 2.**
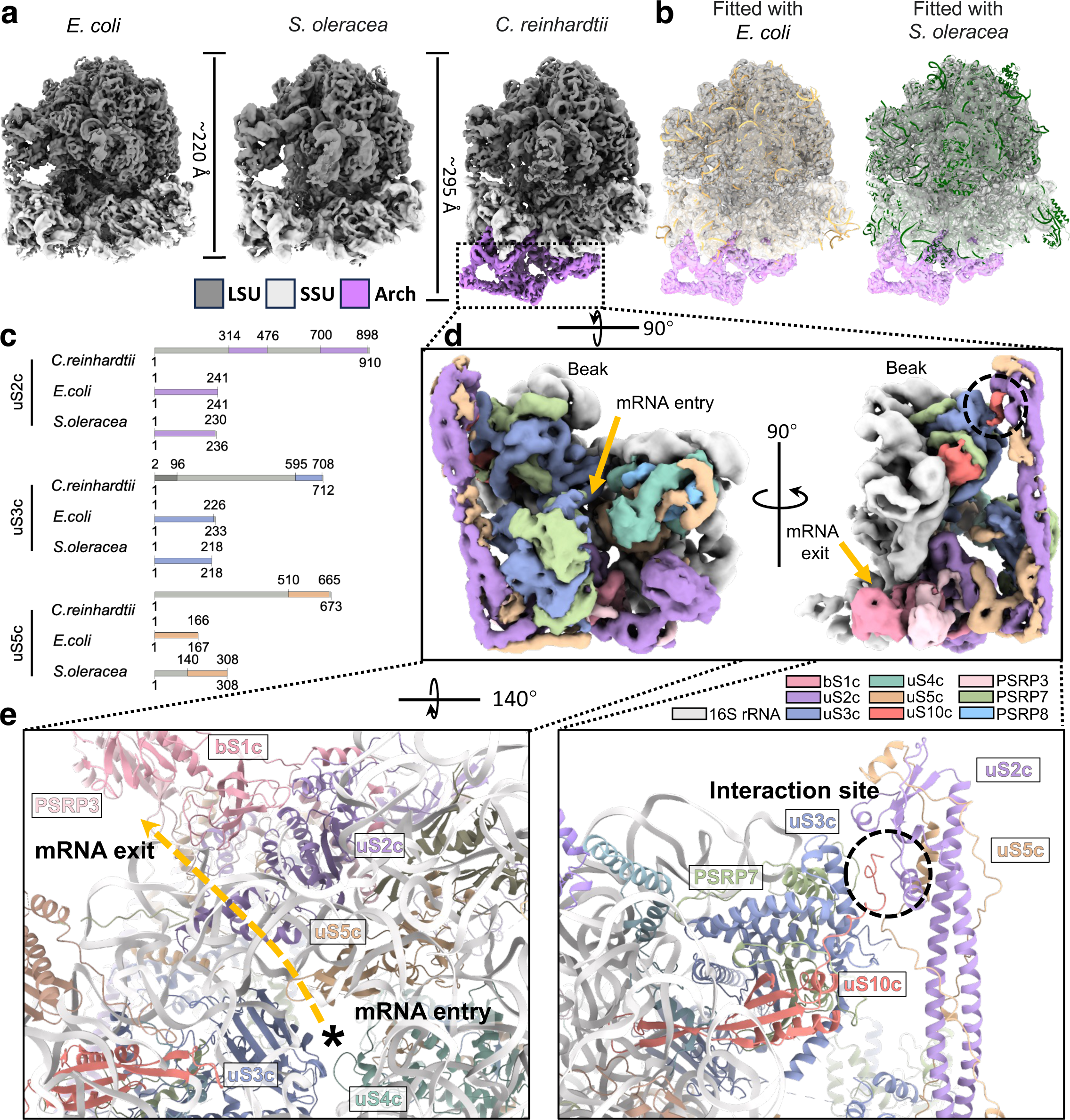
Structure and composition of the arch domain of native chloroplast ribosomes in *C. reinhardtii*. **a,** Structural comparison between ribosomes from *E. coli* (EMD-22607), chloroplast of *S. oleracea* (EMD-3533), and chloroplast of *C. reinhardtii* in this study. For better visualisation, EM density maps are low-pass filtered to 5 Å. **b**, The in-cell structure of chloroplast ribosome in *C. reinhardtii* fitted with atomic models of the ones from *E. coli* (PDB: 4V4A) and *S. oleracea* chloroplast (PDB: 5MMM). **c**, Domain architecture of *C. reinhardtii* uS2c, uS3c and uS5c compared with *S. oleracea* and *E. coli* orthologues, from NCBI CDD annotations. Light grey, insertions and extensions; colour, conserved domains; dark grey, in *C. reinhardtii* uS3c, residues 2-96 return an additional CDD hit (KH-II_30S_S3, cd02412), indicating it was not used to define the insertion boundary discussed here. Residue positions are annotated. **d**, Focused-refined map of the arch density. Constituent proteins are coloured as indicated. The mRNA entry and exit are indicated by golden arrows. The interaction site between the arch and the beak is indicated by the dashed circle. **e**, Zoomed-in model views of the mRNA path and the interaction site between uS10c and uS2c. The mRNA entry is labelled by the asterisk and the mRNA path is demonstrated as the dashed golden line and the uS2c-uS10c interaction site is indicated by the black dashed circle.

Previous proteomic analyses and sequence comparisons identified extensive insertions and extensions in *C. reinhardtii* SSU proteins, particularly uS2c, uS3c and uS5c, together with several chloroplast-specific proteins^12,13^ (Fig. 2c and Supplementary Fig. 2b). Focused refinement resolved the arch to 5.9 Å, revealing secondary-structure elements but precluding unambiguous sequence assignment (Fig. 2d, Supplementary Fig. 1c-e and Supplementary Table 3). We therefore interpreted the density using AlphaFold3^41^ predictions, initially guided by conserved positions in the spinach structure (PDB 5MMM)^17^ and later, where necessary, the recently reported high-resolution single-particle model (PDB 9TVU)^40^.

The initial model adopted an elongated architecture that closely recapitulated the overall shape of the arch-shaped density (Supplementary Fig. 3a, b). The conserved cores of bS1c, uS2c, and uS5c superimposed well on their *S. oleracea* orthologues, with root-mean-square deviations below 1.5 Å (Supplementary Fig. 3c, d). The conserved regions of the three proteins were initially modelled according to their spinach orthologues. These regions included most of bS1c (residues 61-305), the conserved region of uS2c (residues 301-445 and 781-910), and the conserved C-terminal domain of uS5c (residues 486-673). The algal-specific extensions of uS2c and uS5c forming a rod-like assembly composed of uS2c residues 470-758 and uS5c residues 268-470 could be fitted into the STA density with minor adjustment. The remaining regions of bS1c-uS2c-uS5c were manually traced in Coot by rigid-body placement of individual secondary-structure elements.

Using the uS3c structure from the *S. oleracea* chloroplast 70S ribosome as a reference, we confidently modelled the conserved N- and C-terminal regions of uS3c, comprising residues 1-145 and 431-712, respectively (Supplementary Fig. 3e, ii and iii). The internal insertion, comprising residues 146-430 and lacking a counterpart in the spinach protein, was initially left unmodelled.

Because previous proteomic data suggested that PSRP7 and the extended insertion of uS3c are located in the same general region of the 30S subunit, we tested whether these proteins form a complex that accounts for the remaining density. We therefore predicted the uS3c-PSRP7 complex using AlphaFold Server (Supplementary Fig. 3c, right). The previously positioned N- and C-terminal regions of uS3c served as anchors for fitting the predicted complex into the map. This enabled us to trace PSRP7 residues 399-560 in the density (Supplementary Fig. 3e, iii) and uS3c residues 191-430 and PSRP7 residues 166-375 into the density at the turning point of the arch (Supplementary Fig. 3e, iv) with minor manual adjustments and rigid-body fitting in Coot. The final fitted model differed from the AlphaFold prediction by a root-mean-square deviation of ∼2 Å (Supplementary Fig. 3e, iii and iv), indicating that the predicted uS3c-PSRP7 interface closely resembles the conformation observed in the STA map.

An additional density immediately above bS1c could not be assigned unambiguously from our map alone. It may correspond to the unmodelled C-terminal region of bS1c or to another chloroplast ribosome-associated factor, such as PSRP3, as suggested by the proteomic analysis. The assignment of PSRP3 into this density was subsequently clarified by comparison with the recently reported high-resolution SPA cryo-EM structure of the *C. reinhardtii* chloroplast ribosome^40^. Comparison with this structure also enabled PSRP8 to be positioned, which could not initially be modelled independently from our STA map.

Together, this integrative modelling assigns most of the arch to insertions and extensions in uS2c, uS3c and uS5c and to the chloroplast-specific proteins PSRP3, PSRP7 and PSRP8, with bS1c and uS10c contacting the extended region. The arch surrounds both ends of the mRNA channel, consistent with a proposed role in stabilizing or guiding the mRNA trajectory (Fig. 2e, left)^13,40,42^. Its elongated uS2c-uS5c region contacts the beak through a limited uS2c-uS10c interface (Fig. 2e, right), providing a plausible structural basis for arch flexibility.

### Chloroplast ribosomes conduct localized translation on the thylakoid

To examine the native structural heterogeneity of chloroplast ribosomes, we re-extracted particles at 6× binning with their refined coordinates and orientation, followed by global 3D classification. This analysis identified a distinct membrane-bound class comprising 19.7% of the particles, characterized by an additional LSU-associated density extending towards and contacting the thylakoid membrane (Fig. 3a and Supplementary Fig. 4a, b, e). The connecting density is positioned adjacent to the nascent peptide exit, consistent with a role in coupling translation to the thylakoid membrane. Moreover, the membrane-bound chloroplast ribosome structure contains prominent P-site tRNA density (Supplementary Fig. 4b), indicating that these particles represent a translationally engaged state. Mapping the membrane-bound chloroplast ribosomes back into segmented tomograms showed that they were closely associated with thylakoid membranes in their native cellular environment (Fig. 3b). We next compared the local spatial organization of membrane-bound and stromal ribosomes. Their pooled nearest-neighbour (NN) distributions differed for both particle-centre distance and relative Z-axis angular difference (Fig. 3c, d). Membrane-bound particles were shifted towards shorter NN distances and showed a narrower distribution of relative orientations than stromal ones, consistent with the more prominent neighbouring-particle density observed in the membrane-bound chloroplast ribosome (Supplementary Fig. 4a). These results indicate that thylakoid-associated ribosomes adopt a more compact and orientationally constrained local organization at the membrane surface.

**Figure 3.**
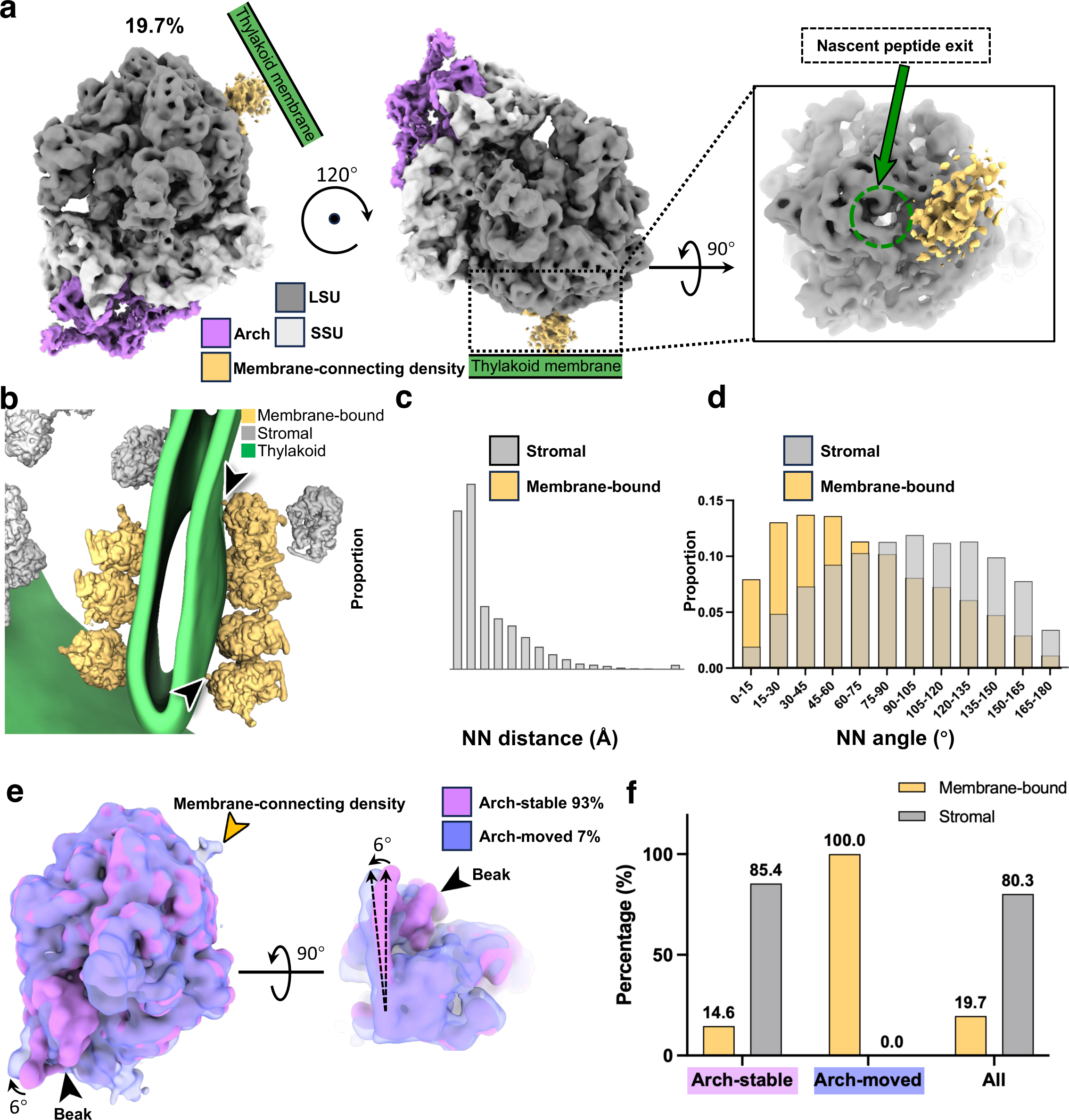
Heterogeneity of native chloroplast ribosomes in *C. reinhardtii*. **a,** STA consensus EM density map of the chloroplast ribosome bound to the thylakoid membrane at 6.9 Å resolution (binning of 1×). Three views are depicted, including a zoomed-in view of the nascent peptide exit on the LSU, which is in proximity to the membrane-connecting density. **b**, A representative mapping back of membrane-bound chloroplast ribosomes together with segmented thylakoid membranes, the contact points are indicated by black arrowheads. **c**, A bar chart displaying the distribution of nearest-neighbour (NN) distance between chloroplast ribosomes in two populations: membrane-bound (gold) and stromal (grey). Chi-square test is applied, *n* of membrane-bound particles = 6,890, *n* of stromal particles = 28,044, *p* < 0.0001. **d**, A bar chart displaying the distribution of nearest-neighbour (NN) angle (Z-axis) between chloroplast ribosomes in two populations: membrane-bound (gold) and stromal (grey). Chi-square test is applied, *n* of membrane-bound particles = 6,890, *n* of stromal particles = 28,044, *p* < 0.0001. **e**, Structural comparison between the arch-stable (93%) and arch-moved (7%) chloroplast ribosomes. For better demonstration, the STA EM density maps are low-pass filtered to 15 Å. The beak and membrane-connecting density are indicated. **f**, A bar chart displaying the proportion of membrane-bound and stromal chloroplast ribosomes in arch-stable, arch-moved, and all chloroplast ribosomes. Fisher’s exact test is applied, *n* of arch-moved particles = 2,430, *n* of arch-stable particles = 32,504, *p* < 0.0001.

Together, these features suggest co-translational targeting of nascent proteins to the thylakoid membrane. At the 6.9 Å resolution of this reconstruction, however, the molecular identity of the membrane connecting density could not be assigned unambiguously. Alb3 is a plausible candidate because of its established function as a thylakoid membrane insertase and recent structural and proteomic evidence linking it to chloroplast ribosomes^15,28,29,40^. Nevertheless, direct identification of the connecting density will require higher-resolution structural data and further biochemical validation.

### Chloroplast ribosomes show prominent arch movement in the membrane-bound state

The arch is connected to the mobile SSU beak through the interaction between uS2c and uS10c, raising the possibility that these regions undergo coordinated conformational changes. To examine this possibility, we performed focused classification centered on the arch and beak domains. This analysis differentiated two populations: an arch-moved class comprising 7% of the particles and an arch-stable class comprising the remaining 93% (Fig. 3e and Supplementary Fig. 4a, c-e). Comparison of these two conformations revealed an approximately 6° displacement of the arch in the arch-moved state. This displacement occurred together with the SSU beak, whereas the LSU densities remained closely aligned between the two classes (Fig. 3e). These observations indicate that the arch and beak move as a coordinated structural unit rather than undergoing independent displacements. Of note, the arch-moved class also contained prominent membrane-connecting density, suggesting a relationship between this conformational state and thylakoid association. We therefore performed an additional classification to determine the membrane-bound fraction within each arch state. Unexpectedly, all particles assigned to the arch-moved class were classified as membrane-bound, whereas only a minority of arch-stable particles showed membrane association (Fig. 3f and Supplementary Fig. 4c, d). The distributions differed significantly between the two conformational populations (Fisher’s exact test, P < 0.0001), demonstrating that arch movement and membrane association are closely-linked. One possible explanation is that membrane engagement constrains the position of the LSU, thereby increasing the relative displacement of the SSU beak and arch or making this displacement more readily detectable by classification. Our analysis establishes an association between the arch movement and membrane-bound state of the chloroplast ribosome. However, higher-resolution structural information of the arch-moved population, together with complementary analysis of their interaction with the membrane, will be required to determine how thylakoid engagement influences chloroplast ribosome conformation.

### Arch components have distinct evolutionary origins but are under purifying selection

To determine whether the distinctive arch is conserved in other species, we conducted phylogenetic analyses of its plastid-encoded components, uS2c and uS3c, and its nuclear-encoded components, uS5c and PSRP7, which differ in their genomic origins and evolutionary histories. A maximum-likelihood phylogeny based on concatenated plastid genes from 23 representatives of Rhodophyta, Chlorophyta and Streptophyta provided a framework for examining their distribution (Fig. 4a and Supplementary Table 4). Within the Chlorophyceae, the uS3c insertion was the most widely distributed, occurring in 93 species spanning most of the sampled families as well as further Chlorophyceae families not shown, including the Sphaeropleales, whereas the uS2c insertion was detected in 27 species, all within the Chlamydomonadales (Fig. 4c, Supplementary Table 6 and Source Data).

**Figure 4.**
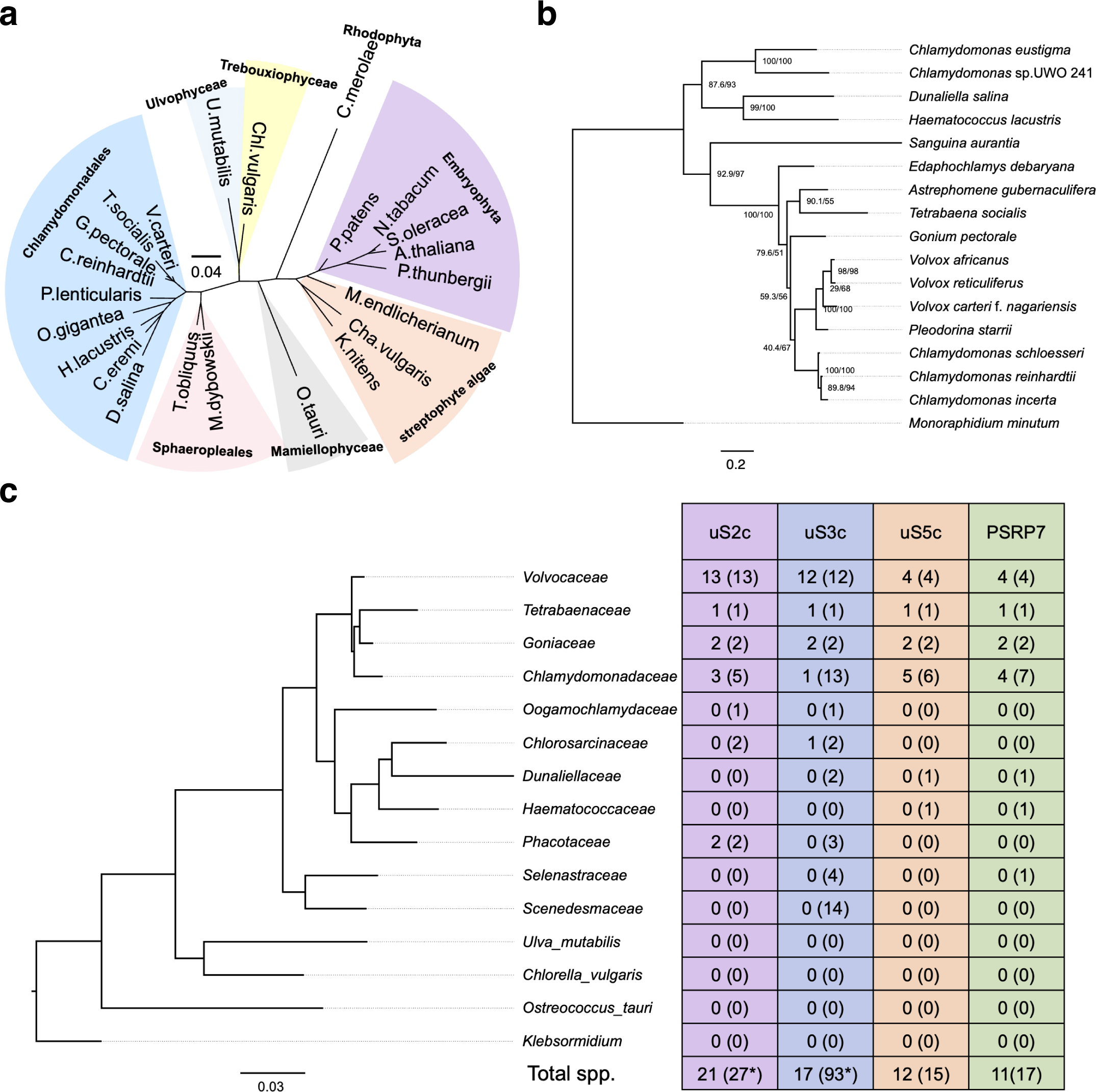
Arch-forming protein components are enriched within the Chlamydomonadales. **a**, Unrooted maximum-likelihood phylogeny inferred from concatenated AtpB, PsaB and RbcL sequences from 23 representatives of Rhodophyta, Chlorophyta and Streptophyta (1,217 amino acids; Q.PFAM+I+R3 model). *Cyanidioschyzon merolae* is included as a distant reference rather than an outgroup. Shaded wedges denote clades, except streptophyte algae, which form a weakly supported grade (9% bootstrap). Scale bar, 0.04 substitutions per site. **b**, Maximum-likelihood phylogeny of PSRP7 homologues from 16 Chlamydomonadales species and *Monoraphidium minutum*, rooted on M. minutum according to the accepted species phylogeny. Node support is shown as SH-aLRT/UFBoot percentages (1,000 replicates); nodes below 80/95 are not interpreted. The sequences of *Chlamydomonas eustigma*, *Chlamydomonas sp.* UWO 241, *Dunaliella salina*, *Haematococcus lacustris* and *Sanguina aurantia* are markedly more divergent than the remaining species, sharing 37.8% mean pairwise identity with them where those species share 73.1% with one another, and are placed basal to the clade they form (100/100); this arrangement follows sequence divergence rather than the accepted species relationships and is not interpreted. Scale bar, 0.2 substitutions per site. **c**, Left, maximum-likelihood phylogeny of representative Chlorophyceae families inferred from concatenated plastid *atpB*, *rbcL* and *psaB* sequences and rooted on *Klebsormidium nitens* (Klebsormidiophyceae). Outgroup counts are shown at class level. Scale bar, 0.03 substitutions per site. Right, numbers of species containing the lineage-specific regions of uS2c, uS3c and uS5c or full-length PSRP7, detected by one-iteration PSI-BLAST searches of NCBI nr. Values are given as *n* (*N*) for ≥40% (≥30%) amino-acid identity, with E ≤ 1 × 10^−20^ and query coverage ≥60%. A value of 0 indicates no hit above these thresholds. uS5c and PSRP7 showed similar distributions, predominantly within the Chlamydomonadales, with PSRP7 additionally detected in the Selenastraceae at the relaxed threshold. Asterisks indicate totals that include species outside the displayed families (uS3c, 39 of 93; uS2c, 1 of 27), all within the Chlorophyceae. Infraspecific taxa were counted separately.

The nuclear-encoded components showed narrower, nearly identical distributions. PSRP7 homologues were recovered from 16 Chlamydomonadales species and the sphaeroplealean *Monoraphidium minutum*, with no homologue detected outside the Chlorophyceae, and all 17 were included in the gene tree. The tree recovered the expected volvocine groups — the three *Chlamydomonas* species and the four Volvocaceae representatives, both at 100/100 — but arranged the five most divergent sequences by their divergence rather than by the accepted species relationships, those sequences sharing 38% mean identity with the remaining species where those share 73% with one another. Neither these placements nor that of *M. minutum* among them is interpreted, and the origin inference below does not depend on them (Fig. 4b and Supplementary Table 5). In the broader homology survey, uS5c and PSRP7 were detected in 15 and 17 species, respectively; the two proteins co-occurred in every identified family except the Selenastraceae, where only PSRP7 was recovered, in *M. minutum*. These results place the detectable origin of PSRP7 at or before the divergence of the Sphaeropleales and Chlamydomonadales and that of uS5c within the Chlamydomonadales, supporting stepwise recruitment of nuclear-encoded proteins onto an older plastid-encoded scaffold.

To assess functional constraint, codon alignments of the four proteins were retrieved independently using full-length queries and analysed for selection in HyPhy, after excluding sequences whose length was inconsistent with that of the remaining taxa (*Edaphochlamys debaryana* from uS2c and uS3c; *Sanguina aurantia* from uS5c). Because these retrievals used full-length queries rather than the lineage-specific regions surveyed in Fig. 4c, their sample sizes are not directly comparable with it, except for PSRP7, for which the same query was used in both. All four proteins showed low gene-wide dN/dS values (ω = 0.115-0.141). FEL identified significant purifying selection at 48-60% of codons, roughly five-to six-fold more than expected by chance at this threshold, and no codon under positive selection in any of the four alignments. At the gene level, no BUSTED test of episodic diversifying selection remained significant after Holm-Bonferroni correction across the four genes (Supplementary Table 7 and see Methods). Thus, despite their different genomic origins and evolutionary ages, both plastid- and nuclear-encoded arch components are predominantly constrained by purifying selection, supporting the arch as a functionally maintained structural feature.

## Discussion

Here we determined the native structure, conformational heterogeneity and spatial organisation of the *C. reinhardtii* chloroplast ribosome in intact cells. The 4.9 Å consensus reconstruction, with local resolution reaching 4.4 Å, revealed a prominent arch-shaped extension of the SSU. Integrative modelling of the focused arch reconstruction identified sequence insertions and extensions in uS2c, uS3c, and uS5c, together with the chloroplast-specific proteins PSRP3, PSRP7 and PSRP8, as its principal components. Classification further resolved thylakoid-associated ribosomes and an arch-moved state in which the arch and SSU beak were displaced together. The strong enrichment of this state among membrane-bound particles, combined with their distinct nearest-neighbour distance and orientation distributions, links chloroplast ribosome conformation to membrane association and localized translation in cells.

The position of the arch around the mRNA entry and exit channels suggests that it may influence chloroplast mRNA translation. The unusually large forms of uS2c, uS3c and uS5c, and the presence of PSRP7, were first identified by proteomic analyses of the *C. reinhardtii* chloroplast ribosome^12,13^. Our structure places these extensions within a continuous SSU domain and shows that uS3c, uS4c and uS5c surround the mRNA entry region, whereas bS1c and PSRP3 lie near the exit. PSRP7 is particularly of note because it contains S1 RNA-binding domains and is produced as part of a polyprotein containing translation elongation factor Ts domains^42^. Plastid-specific ribosomal proteins can be required for ribosome accumulation and chloroplast translation^22^, but the precise contributions of the individual arch components were not known. The structure therefore supports a role of arch subunits in mRNA recruitment or stabilisation, rather than establishing a specific regulatory mechanism.

The RNA-binding properties of PSRP7 also raise the possibility that the arch participates in coordinating transcription and translation. In bacteria, these processes can be physically coupled within an expressome through contacts between RNA polymerase, the ribosome and the transcription factors NusG and NusA^43–46^ (Supplementary Fig. 5a). A chloroplast NusG homologue in *Arabidopsis* associates with the plastid-encoded RNA polymerase and interacts with the ribosomal proteins uS5c and uS10c, providing evidence for a related coupling mechanism in plastids^47^. Transcriptionally active nucleoids and chloroplast translation zones are also spatially coordinated in *C. reinhardtii*^6,30^. Given its S1 domains and position near the mRNA channel (Supplementary Fig. 5b), PSRP7 could help retain nascent transcripts or recruit RNA-associated factors to the ribosome. However, our maps do not resolve RNA polymerase or a direct interaction between PSRP7 and the transcription machinery. PSRP7 should therefore not yet be considered as a chloroplast equivalent of NusG or NusA. It would be interesting to test whether it contributes to an expressome-like assembly via direct biochemical and structural evidence in the future.

Our analysis also provides insight into thylakoid-localized translation. The additional density connecting the ribosome to the membrane lies adjacent to the nascent peptide exit and the membrane-bound ribosomes are resolved with prominent P-site tRNA density, consistent with a translating ribosome engaged in co-translational membrane targeting. Alb3, a member of the Oxa1/YidC/Alb3 protein family, whose mitochondrial members can mediate ribosome tethering^48,49^, is therefore a plausible component of the connecting density in the chloroplast. Alb3 mediates co-translational insertion of plastid-encoded thylakoid proteins, interacts directly with chloroplast ribosomes through its stromal C-terminal region^29^, and has been reported in the chloroplast ribosome interactome^15^. Nevertheless, the connecting density cannot be assigned unambiguously at the present resolution and may contain contributions from Alb3, the nascent peptide, chloroplast signal-recognition-particle components or additional membrane-biogenesis factors. The density should therefore be described as Alb3-compatible here and requires further validation in the future.

The coupling between arch movement and membrane association provides a further view of chloroplast ribosome dynamics. In the arch-moved class, the arch and beak undergo an approximately 6° coordinated displacement while the LSU remains aligned. All particles assigned to this class were membrane-bound, whereas only a minority of arch-stable particles were associated with the thylakoid. One interpretation is that membrane tethering restricts LSU motion and thereby makes relative movement of the SSU and arch more readily detectable. Alternatively, engagement of a translating ribosome with the thylakoid insertion machinery could alter the conformation of the SSU through the limited uS2c-uS10c interface connecting the arch to the beak. These possibilities are not mutually exclusive. The shorter nearest-neighbour distances and narrower relative-orientation distribution of membrane-bound particles further suggest that the thylakoid imposes spatial constraints on translating ribosomes, potentially favouring ordered translation assemblies. Of note, the associations observed in our study do not establish whether membrane binding induces arch movement or whether this movement plays a direct role in translation.

Our evolutionary analyses indicate that the arch is an evolutionary mosaic assembled through the stepwise recruitment of proteins with distinct genomic origins and evolutionary histories. The plastid-encoded uS3c insertion is broadly distributed across the Chlorophyceae, whereas the uS2c insertion is restricted to the Chlamydomonadales. Among the nuclear-encoded components, the detectable distribution of PSRP7 places its origin at or before the divergence of the Sphaeropleales and Chlamydomonadales, whereas the extended uS5c region appears to have emerged later within the Chlamydomonadales. The similar distributions of PSRP7 and the uS5c extension within this lineage suggest their coordinated recruitment or retention.

Despite their different evolutionary ages, all four major arch components are predominantly constrained by purifying selection, with no codon under positive selection and no gene-wide evidence of episodic diversifying selection after correction across the four genes. Although evolutionary conservation alone does not establish function, these selective constraints support the arch as a functionally maintained feature rather than a neutral structural elaboration.

Recently, Waltz and colleagues independently reported the *C. reinhardtii* chloroplast ribosome using *in situ* SPA cryo-EM and cryo-ET^40^. High-resolution SPA enabled an essentially complete molecular model, establishing the composition of the domain termed the SSU arm in their study and the arch here. They also observed flexibly tethered, thylakoid-associated ribosomes and proposed that the SSU extension stabilises the mRNA path and contributes to polysome organisation. These findings provide definite molecular assignments that complement and strengthen our integrative cryo-ET study. In turn, our analysis extends this in-cell description by resolving coordinated arch and beak movement, demonstrating its association with the membrane-bound state, quantitatively comparing the local organisation of membrane-bound and stromal particles, and defining the phylogenetic distribution and selective constraints of the arch components. Together, the two studies establish the arch as a genuine feature of the native *C. reinhardtii* chloroplast ribosome while providing complementary molecular and cellular perspectives. Notably, in-cell reconstructions from both studies resolved the arch in all chloroplast ribosomes, whereas only a small fraction (∼24%) of the purified particles retained resolvable arch density in the related study^40^, underscoring the ability of cryo-ET and STA to capture native molecular structures while avoiding perturbations associated with biochemical isolation.

Future studies would therefore combine targeted genetics, functional measurements and higher-resolution in-cell imaging. Because the conserved cores of uS2c, uS3c and uS5c are likely required for ribosome assembly, selective truncation of their lineage-specific insertions or extensions would be more informative than complete gene deletion. Complemented or inducible mutants should be assessed for ribosome assembly, polysome formation, chloroplast translation, transcript-specific ribosome occupancy, photosynthetic performance and acclimation to changing light conditions. Structural analysis of these mutants could determine whether individual extensions are required for arch assembly, mRNA positioning, and transcription-translation coupling.

Together, our findings connect a lineage-specific ribosomal extension to chloroplast ribosome conformation, thylakoid association, spatial organisation and evolutionary diversification. Moreover, they illustrate how in-cell structural analysis can integrate molecular structure with cellular context, dynamics and evolution, providing a powerful exploratory approach not only for structural biology but also for cell biology, organelle genetics and evolutionary research.

## Methods

### Cell culture and cryo-sample preparation

Wild-type *C. reinhardtii* CC-4533 cells obtained from obtained from the Chlamydomonas Resource Center (https://www.chlamycollection.org/, funded by the US National Science Foundation) were maintained in Tris-acetate-phosphate medium under a 12 h light/12 h dark cycle at room temperature, with illumination at 12,000 lux and shaking at 150 rpm. Cells were harvested 6 h after the start of the light phase. Approximately 3.5 µl of cell suspension at 1.5 × 10⁶ cells ml^-^¹ was applied to a glow-discharged Quantifoil R2/1, 300-mesh holey-carbon grid, blotted from the reverse side for 8 s and plunge-frozen in liquid ethane.

### Cryo-FIB milling

Cryo-FIB milling was performed on an Aquilos 2 cryo-FIB/SEM operated at 30 kV with a gallium ion beam. Organometallic platinum was deposited for 30 s using the gas-injection system before milling. Milling currents were 0.1-0.5 nA, and final polishing currents were 30-50 pA. No sputter coating was applied before or after polishing. In total, 64 lamellae were prepared (Supplementary Table 1).

### Cryo-ET data collection

Tilt series were acquired on FEI Titan Krios G3 microscopes operated at 300 kV and equipped with a Falcon 4i direct detector and a Selectris X energy filter with a 10 eV slit. Data were collected without super-resolution at a physical pixel size of 1.903 Å per pixel. The defocus range was −2 to −5 µm in 0.25 µm increments. A dose-symmetric acquisition scheme with a 54° tilt span, 2° increments and groups of three tilts was used. The total dose was 137.5 e^-^ Å^-^², with 10 movie frames recorded per tilt. The dataset comprised 64 lamellae and 260 tomograms at 1.903 Å per pixel (Supplementary Table 2).

### Alignment of tilt series and tomogram reconstruction

The frames of each tilt series were corrected for beam-induced motion using MotionCor2^50^. The gain correction was performed in parallel with the motion correction run by a home-brewed script. New stacks were generated and aligned using IMOD^51^ version 4.11.1 by patch tracking, and tomograms were reconstructed at binning of 6× with a pixel size of 11.418 Å/pixel.

### Template matching

A 40 Å low-pass-filtered chloroplast ribosome template adapted from EMD-3533 was used for initial template matching in emClarity 1.5.0.2^52^ at 6× binning. A deliberately large particle-picking threshold enabled an exhaustive search within selected tomographic regions. Candidate particles were inspected manually in ChimeraX^53^, and false positives, predominantly cytoplasmic ribosomes, were removed (Supplementary Fig. 1a).

### Subtomogram averaging

#### Refinement of the consensus map

The manually cleaned set of 49,911 particles was imported into RELION 4.0^54^. Iterative 3D classification with alignment at 6× binning was followed by stepwise refinement from 6× to 1× binning (Supplementary Fig. 1a). Classes with ribosome-like morphology were retained, yielding 34,934 particles for the consensus reconstruction. Refinement with C1 symmetry produced a 4.9 Å global map, with local resolution reaching 4.4 Å and no evident preferred orientation. Resolution was assessed using the gold-standard Fourier shell correlation (FSC) 0.143 criterion (Supplementary Fig. 1a, b, d, e and Supplementary Table 3).

### Focused arch refinement

For focused refinement, a mask encompassing the arch density and part of the SSU was applied to the consensus particle set. Refinement at 1× binning produced a 5.9 Å arch map without evident preferred orientation (Supplementary Fig. 1c-e and Supplementary Table 3).

### Structure prediction and model building of the arch

Protein structures and protein complexes were predicted using AlphaFold Server, implementing AlphaFold3^41^, with default settings. Predictions were performed before the high-resolution in vitro structure of the *C. reinhardtii* chloroplast ribosome become available^40^. Five models were generated for each prediction, and the model with the highest-ranking score was selected for subsequent analysis.

Predicted models were initially placed into the STA density map by rigid-body fitting in UCSF Chimera v1.18^55^. Models were then manually adjusted and subjected to rigid-body fitting in Coot v0.9^56^. Structural superpositions and root-mean-square-deviation calculations were performed using PyMOL v3.1.6.1. Molecular graphics were prepared using PyMOL v3.1.6.1 (Schrödinger) and UCSF ChimeraX v1.12^53^.

### Classification

For the membrane-bound subset, consensus particles were re-extracted at 6× binning and classified without alignment. Selected particles were refined stepwise from 6× to 1× binning, producing a 6.9 Å map from 6,890 particles (Supplementary Fig. 4a, b, e and Supplementary Table 3). For analysis of arch states, a mask was placed over the arch density and classification without alignment at 6× binning identified 2,430 arch-moved particles and 32,504 arch-stable particles (Supplementary Fig. 4a, c-e and Supplementary Table 3). Stepwise refinement yielded maps at 8.4 Å and 5.2 Å, respectively. Particles from each arch state were then re-extracted at 6× binning and subjected to 3D classification without alignment to identify membrane-bound subclasses.

### Segmentation and mapping back

Membranes were initially segmented in tomograms using MemBrain v2^57^ and then imported into ChimeraX for manual cleaning and refinement. Ribosome particles were mapped back into the tomographic volumes using their refined coordinates and orientations. For visualization, ribosome models in segmented volumes were generated from low-pass-filtered structures obtained in this study. Visualizations were prepared in ChimeraX and ArtiaX^58^.

### Nearest-neighbour (NN) analysis

Particles were grouped by tomogram, and the nearest neighbour of each particle was defined as the closest other particle within the same tomogram. Coordinate pairs separated by less than 70 Å were treated as duplicate picks and removed before subsequent analysis. Nearest-neighbour distance was measured between particle centres. The nearest-neighbour Z-axis angle was calculated from the local Z directions defined by the particle orientations. Binned distributions were compared using chi-square tests and plotted in Prism 10.

### Phylogenetic and selection analyses

#### Homologue identification and distribution survey

Domain boundaries in *C. reinhardtii* uS2c (ASF83566.1), uS3c (ASF83570.1) and uS5c (AAM18796.1) were defined by comparison with *S. oleracea* and *E. coli* K-12 orthologues in the NCBI Conserved Domain Database using default parameters. In uS2c the rps2 domain (CHL00067) was recovered in two segments flanking an intervening region absent from both orthologues; in uS3c, KH-II_30S_S3 (cd02412) and RpsC (COG0092) flank a chlorophyte-specific insertion; in uS5c, RpsE (COG0098) is preceded by an N-terminal extension. To survey the distribution of these lineage-specific regions, uS2c residues 476-700, uS3c residues 97-595, uS5c residues 1-510 and full-length PSRP7 (AAU93599.1) were used as queries for PSI-BLAST (one iteration) against the NCBI non-redundant protein database.

Hits were retained at E ≤ 1 × 10−20 and query coverage ≥ 60%, restricted to the Chlorophyta and counted at two identity thresholds, ≥ 40% and ≥ 30%. Multiple accessions of the same species were counted once, infraspecific taxa were counted separately, and species were assigned to families following AlgaeBase. All retained hits, with their identity, query coverage, E value and the reason for each exclusion, are provided in Source Data.

### Phylogenetic analyses

Coding sequences for *atpB*, *psaB*, and *rbcL* were extracted from complete plastid genomes, translated, aligned separately with MAFFT v7.526^59^ (L-INS-i) and concatenated. For the species phylogeny, 23 representatives of the Rhodophyta, Chlorophyta and Streptophyta gave 1,217 amino-acid positions, and a maximum-likelihood tree was inferred with IQ-TREE v3.0.1^60^ under the Q.PFAM+I+R3 model selected by ModelFinder^61^ (BIC), with 1,000 ultrafast bootstrap replicates and SH-aLRT. The tree is shown unrooted, with Cyanidioschyzon merolae included as a distant reference rather than an outgroup. The family-level Chlorophyceae cladogram was inferred from the same genes using one exemplar species per family, or one per class for the four outgroup lineages, and rooted on the Klebsormidiophyceae exemplar *Klebsormidium nitens*. For the PSRP7 gene tree, homologues were retrieved by PSI-BLAST as above at ≥ 30% identity, sequences annotated as PSRP7– EF-Ts polyproteins were trimmed to the PSRP7 domain, corresponding to residues 1–560 of the C. reinhardtii protein, and all 17 species in which a homologue was detected were included. Sequences were aligned with MAFFT v7.526 (--auto) and ambiguously aligned columns were removed with trimAl v1.5 (-automated1), retaining 484 of 842 columns (32 constant, 347 parsimony-informative, 470 distinct site patterns), and a tree was inferred with IQ-TREE v3.0.1 under the LG+F+G4 model selected by ModelFinder (-m MFP) under BIC, with 1,000 ultrafast bootstrap and 1,000 SH-aLRT replicates. The tree was rooted on *Monoraphidium minutum* following the accepted sister relationship of the Sphaeropleales and Chlamydomonadales, so the rooting derives from the species phylogeny rather than from the PSRP7 alignment; nodes with SH-aLRT < 80 or UFBoot < 95 were not interpreted.

Sequences used for the species phylogeny and the PSRP7 phylogeny are listed in Supplementary Table 4 and 5, respectively, and the exemplar species used to generate the family-level cladogram in Supplementary Table 6.

### Selection analyses

Sequences for selection analysis were retrieved separately using the full-length *C. reinhardtii* proteins as queries at ≥ 40% identity under the same E-value and coverage criteria. Sequences whose length was inconsistent with that of the remaining taxa were excluded: *Edaphochlamys debaryana* from uS2c and uS3c, and *Sanguina aurantia* from uS5c. Codon alignments were generated from the protein alignments and corresponding coding sequences with PAL2NAL v14 (-nogap), using genetic code table 11 for the plastid-encoded uS2c and uS3c and table 1 for the nuclear-encoded uS5c and PSRP7. Analyses were performed in HyPhy v2.5.98^62^ via the Datamonkey server with site-to-site synonymous rate variation enabled throughout. Each alignment was first screened for recombination with GARD; the breakpoints recovered involved the same short-internode volvocine lineages across genes and yielded partitions too short to identify topologies reliably, so they were attributed to phylogenetic uncertainty rather than recombination and whole-alignment analyses were used throughout. Gene-wide episodic diversifying selection was tested with BUSTED and assessed against a Holm-Bonferroni correction for four tests at a family-wise α of 0.05. Site-level selection was assessed with FEL, with sites reported at p ≤ 0.1; no correction across codons was applied, and observed counts were instead compared with the number expected by chance at this threshold.

## Data availability

All data required to evaluate the conclusions are provided in the Article and its Supplementary Information. The subtomogram-averaged maps are deposited in the Electron Microscopy Data Bank (EMDB) under the accession codes: EMD-59302, −59303, −59304, −59305, and −59306 for the consensus map of in-cell structure of chloroplast ribosome, the arch domain, the arch-moved class, the arch-stable class, and the membrane-bound class, respectively. Source data are provided with this paper.

## Acknowledgements

We thank Dr. L. Carrique, Dr. H. Duyvesteyn and Dr. J. Gilchrist for support with data collection, and Dr. J. Sun for helpful discussions, we also thank Dr. Emma F Harding for help in phylogenetic analysis. Y.S. was supported by a Canadian Institutes of Health Research fellowship (194032) and an EMBO fellowship (ALTF 96-2024). We acknowledge the Oxford Particle Imaging Centre for access to the Krios G3 cryo-electron microscope and Diamond Light Source for access to, and support from, the UK national Electron Bio-imaging Centre (eBIC; proposal NT29812). Computation was performed at Diamond Light Source with support from the Wellcome Trust Core Award (203141/Z/16/Z) and the NIHR Oxford Biomedical Research Centre. This work was supported by US National Institutes of Health grants U54AI170791 and R21AI184080 (P.Z.); Wellcome Investigator Award 206422/Z/17/Z and Wellcome Discovery Award 311427/Z/24/Z (P.Z.); European Research Council Advanced Grant 101021133 (P.Z.); and the Chinese Academy of Medical Sciences Innovation Fund for Medical Science (2024-I2M-2-001-1; P.Z.); European Research Council (ERC) [101001623-PALVIREVOL to A.K.]. The funders had no role in study design, data collection and analysis, the decision to publish or preparation of the paper.

## Author contributions

P.Z. and Z.H. conceptualized this study. Z.H., Y.S. and Z.Z. designed the experiments. Z.H. prepared samples with assistance from P.L., performed cryo-FIB milling, collected the cryo-ET data and reconstructed the tomograms. Z.H. performed STA, segmentation and classification analyses. Y.S. performed structural prediction and modelling. Z.Z. performed the phylogenetic analysis under A.K.’s supervision. Z.H., Y.S. and Z.Z. prepared the figures. Z.H., Y.S., Z.Z. and P.Z. wrote the paper with input from all authors.

